# Transient cortical activity modulates somatosensory gating during postural control

**DOI:** 10.64898/2026.10.20.760602

**Authors:** Kimiya Fujio, Hiroki Obata, Nadaka Hakariya, Noritaka Kawashima

## Abstract

Human somatosensory responses vary with movement-related gating as well as with fluctuations in ongoing cortical activity. During standing, both somatosensory responses and cortical activity change in response to postural demands. However, little is known regarding whether moment-to-moment fluctuations in cortical activity contribute to variation in somatosensory responses during different postural tasks. In this study, we investigated whether somatosensory-evoked potentials (SEPs) during postural control are associated with prestimulus cortical activity, focusing on oscillatory amplitudes and corticocortical connectivity. SEPs evoked by tibial nerve stimulation were recorded in 18 healthy adults under four postural conditions. Canonical correlation analysis (CCA) was used to extract single-trial SEP amplitudes from multichannel electroencephalography (EEG). The results showed that the early component of CCA-SEPs, N80, was significantly reduced according to postural demands. A higher prestimulus amplitude in the β-frequency band in the central cortical regions was associated with a larger N80 amplitude. Moreover, modulation of α-band corticocortical connectivity involving frontal and parietal regions covaried with N80 reduction. Clustering analysis using β-amplitude and α-connectivity as features further indicated that multivariate prestimulus cortical activity accounted for variability in N80 amplitude. These findings revealed that somatosensory responses vary through posture-related somatosensory gating and as a function of ongoing fluctuations in cortical activity. Clarifying the temporal dynamics of somatosensory processing will help elucidate the underlying physiological mechanisms of standing balance and contribute to the development of novel physiological assessments of postural impairments.

## Introduction

Stable human posture is achieved through continuous visual, vestibular, and somatosensory inputs that are integrated and weighted according to the environment and task demands. The somatosensory system provides the central nervous system with information regarding body configuration and surface conditions to maintain stable postural balance. Somatosensory-evoked potentials (SEPs) and somatosensory-evoked magnetic fields (SEF) are widely used for an electrophysiological assessment of somatosensory processing. It has been established that the SEP/SEF are generally attenuated during voluntary movements, such as tonic muscle contraction, finger tapping, and ankle flexion (Starr & Cohen, 1985; Kakigi *et al*., 1995; Morita *et al*., 1998; Lin *et al*., 2000). This phenomenon is described as movement-related somatosensory gating, involving multiple peripheral and central processes. Somatosensory gating is considered to suppress sensory inputs generated by self-movement, thereby facilitating the processing of task-relevant somatosensory information.

In addition to voluntary movements, somatosensory processing is influenced by transient cortical activity. Such transient fluctuations in cortical activity have been linked to moment-to-moment variability in neural responses across several sensory modalities, including visual (Becker *et al*., 2008; Iemi *et al*., 2019), auditory (Romani *et al*., 1988), and somatosensory stimuli (Nikouline *et al*., 2000; Stephani *et al*., 2020). In the somatosensory domain, cortical oscillatory amplitudes in the α- or β-frequency bands (8–13 Hz and 13–30 Hz, respectively) covaried with the early component of evoked responses (Nikouline *et al*., 2000; Cebolla *et al*., 2009; Forschack *et al*., 2017; Stephani *et al*., 2020), as well as tactile perception (van Ede *et al*., 2010; Baumgarten *et al*., 2016). During rest, lower prestimulus α-power centred on sensorimotor regions predicts larger SEPs to somatosensory stimuli (Stephani *et al*., 2020), although another study reported only a weak overall positive association (Nikouline *et al*., 2000). In a stimulus detection task, larger SEPs were observed in perceived trials, and a nonlinear relationship was observed between μ-power over the somatosensory cortex and early SEP amplitude (Zhang & Ding, 2010). By contrast, in a tactile simultaneity judgement task, prestimulus β-power was associated with SEF responses over primary and secondary somatosensory areas in opposite directions, showing positive and negative associations, respectively (Lange *et al*., 2012), whereas the relationship between α-power and SEF responses depends on the stimulus onset asynchronies. Evidence indicates that somatosensory responses, tactile perception, and subsequent behavioural performance inherently fluctuate despite identical sensory inputs, partly as a function of ongoing cortical activity (Arieli *et al*., 1996). These findings suggest top-down control of sensory processing, which is attributed to the cortical excitability and network state preceding sensory stimulation. Importantly, this relationship is not fixed, but can vary with task-dependent changes in the baseline cortical state.

When maintaining posture, cortical activity, including top-down signals, varies flexibly according to postural demands. For example, the oscillatory power at the α-frequency band in the central region decreased during more challenging tasks, such as tandem stance or standing with a cognitive task (Edwards *et al*., 2018; Kahya *et al*., 2022). Cortical β-power also increases in the frontal and parietal regions in young adults, but decreases in aged adults, when standing on a foam surface or in a tandem position (Malcolm *et al*., 2021; Hu *et al*., 2023). Cortical connectivity was further reorganised according to sensory information during standing. In a single-leg stance on an unstable surface, α-cortical connectivity extends from the occipital to the parietal and parieto-central areas (Mierau *et al*., 2017). Visual conditions also affect cortical connectivity in the frontal area, which is more dominant in the eyes-open condition, whereas temporo-parieto-central networks are enhanced in the eyes-closed condition (Barollo *et al*., 2022). Both cortical oscillatory power and connectivity are flexibly modulated according to the postural demands and stability requirements of the postural task.

Posture-related somatosensory gating may be influenced by both posture-dependent changes in cortical activity and moment-to-moment fluctuations in ongoing cortical activity. However, it remains unclear whether the relationship between prestimulus cortical activity and SEPs varies across different cortical states associated with postural tasks. Clarifying this question will improve our understanding of the neural mechanisms underlying the dynamic regulation of somatosensory processes for standing balance and may contribute to the development of electrophysiological assessments for individuals with postural disorders. In this study, we aimed to determine whether ongoing cortical activity is associated with somatosensory gating during postural control. To address this question, we examined the relationship between oscillatory cortical activity immediately before stimulation and lower-limb SEP amplitude at the single-trial level. Because cortical oscillatory activity fluctuates over time during motor behaviour, even during tonic motor output or postural maintenance (Echeverria-Altuna *et al*., 2022; Fujio *et al*., 2024), we hypothesised that prestimulus cortical activity during postural control is associated with trial-by-trial variability in SEP amplitude. The analysis focused on an early component of lower-limb SEP peaking at approximately 80 ms (N80), which exhibited a relatively high signal-to-noise ratio (SNR). First, we examined whether fluctuations in cortical α-and β-amplitudes immediately before electrical stimulation were associated with the lower-limb SEP amplitude. Across different postural conditions, prestimulus α- and β-amplitude and SEP amplitude were calculated on a trial-by-trial basis, accounting for within-participant variability (that is, single-trial analysis). Second, we investigated whether prestimulus cortical connectivity shaped by postural demands specific to each condition was associated with SEP amplitude. Focusing on cortical regions presumed to underlie SEP modulation, a condition-averaged analysis was performed to determine whether prestimulus connectivity patterns were associated with SEP modulation during postural control. Finally, we performed a clustering analysis to examine whether multivariate cortical features could account for SEP modulation. Using β-amplitude and α-connectivity as clustering features, trial-by-trial prestimulus neural activities were classified into a small number of neural states, and SEP amplitudes were compared across the identified states.

## Methods

### Ethical approval

The experimental procedure was approved by the Ethics Committee of the Research Institute of the National Rehabilitation Centre for Persons with Disabilities (reference number 2026-014). All procedures adhered to the principles outlined in the Declaration of Helsinki. All participants were informed of the study purpose and any potential risks and provided written informed consent before participation.

### Participants

Eighteen healthy adults participated in this study (26.6 ± 1.5 years, 164.6 ± 1.8cm, 11 male). Data from one male participant were excluded because of artifacts presumed to be caused by sweating. Finally, a dataset from 17 participants was used for the statistical analysis. The exclusion criteria were history of neurological disease, motor impairment, or peripheral nerve impairment.

### Data acquisition

The electroencephalography (EEG) signals were recorded at 2,048 Hz using a 64-electrode system (ActiveTwo system^®^, Biosemi, the Netherlands). The electrodes were placed in accordance with the international 10–10 system. A manufacturer-recommended CMS/DRL reference configuration over the occipital region was employed. Surface electromyographic (EMG) activities were measured from five muscles in the left lower limb, including the tibialis anterior, soleus (SOL), medial gastrocnemius, rectus femoris, and biceps femoris muscles (MEG-6109 bioelectric amplifier^®^, Nihon Kohden, Japan, sampling rate: 10 kHz, bandpass filter: 15–3,000 Hz). Bipolar electrodes (Ag-AgCl, 7 mm diameter) were attached 1.5 cm apart on the muscle belly and wrapped with a thin elastic bandage after cleaning with alcohol. To capture postural sway, the ground reaction forces were recorded using a single force plate (EFP-S-1.5KNSA13^®^, Kyowa, Japan, sampling rate: 10 kHz). Electrical stimulation was delivered to evoke SEPs using a constant-current stimulation system (SEN-7203 and SS-104J^®^, Nihon Kohden, Japan). The timing of electrical stimulation and monitoring of EMG activation levels were controlled using LabVIEW software (National Instruments, USA).

### Experimental procedures and tasks

Four experimental conditions were performed with recording of EEG, EMG, and ground reaction forces: sitting on a stool (SIT), quiet-normal standing (QS), sitting on a stool with SOL contraction (CO), and standing on a single wooden block (WD; height: 7.5 cm, length: 42.0 cm, width: 7.5 or 6.0 cm). The SIT condition served as a reference condition representing a resting posture. In the QS condition, the feet were positioned parallel to each other, with a distance of 15 cm between the centres of the heel. In the CO condition, participants were required to maintain SOL muscle activity at a level comparable to that during normal standing (that is, QS) and avoid ankle and knee movements as much as possible. Visual feedback for the target activity level was provided in the practice session only and was removed during the recording. In the WD condition, one of the two wooden blocks was selected according to the foot length of the participant and placed on the force plate. The foot position was the same as that in the QS condition, and it was set so that the metatarsal heads were not placed on the edge of the wood. The QS and WD conditions differed in the restriction of the base-of-support in the anteroposterior direction.

Participants were instructed to maintain their gaze on a visual target 1.5 m in front of them and to avoid deliberate movements, such as shaking their heads or hands. Before the experiment, participants practiced maintaining the required postures. The order of the experimental conditions was randomised, and each condition was performed in four blocks. SEPs were recorded for 50 stimulations per block.

### Peripheral nerve stimulation for SEP

Electrical stimulation was delivered to the distal tibial nerve at the posterior medial malleolus. The optimal stimulation site was first identified using bipolar surface electrodes with reference to hallux flexion. The electrodes were replaced with disposable electrodes. To obtain clearer single-trial SEPs, the electrical stimulation consisted of a train of five rectangular pulses (pulse duration: 1.0 ms, inter-pulse interval: 3.0 ms), which is commonly used to elicit cutaneous reflexes (Nakajima *et al*., 2008). Stimulus intensity was set at 1.3 × the radiating threshold, defined as the lowest intensity at which participants perceived the radiation of the stimulus to the sole. To avoid the activation of nociceptive pathways, the stimulus intensity was confirmed not to induce pain in each participant.

### Data Analysis

#### Postural sway and muscle activities

To estimate standing balance during the two standing conditions, QS and WD, the centre-of-pressure (COP) was computed as an indicator of postural sway. The force data were filtered using a fourth-order, zero-phase-lag Butterworth low-pass filter with a 10-Hz cutoff frequency. Using the vertical ground reaction forces and moments in the AP and mediolateral directions, the time series of the COP was calculated, and COP velocity (COPv) was obtained by temporal differentiation as follows:

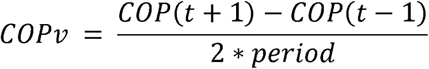

where *t* is the time point, and *period* is the sampling interval.

To confirm the required performance in the CO condition, SOL activity was quantified for 200 ms during the prestimulus period. A Butterworth band-pass filter (15–450 Hz, 4^th^ order) was applied, and the root-mean-square was calculated in every trial under all conditions (SOL-RMS). Both variables were standardised into z-scores across the analysed conditions for each participant.

#### EEG preprocessing

The EEG time series were preprocessed using EEGLAB (Delorme & Makeig, 2004), an open-source toolbox running in MATLAB (MathWorks Inc., USA). Figure 1 outlines the EEG processing. The EEG data were band-pass filtered using a zero-phase filter with a low cutoff of 1.0 Hz and a high cutoff of 300 Hz and then downsampled to 2,000 Hz. After reducing line noise using the cleanLineNoise function in the EEGLAB plugin, bad channels with obvious artifacts were removed via visual inspection.

**Figure 1.**
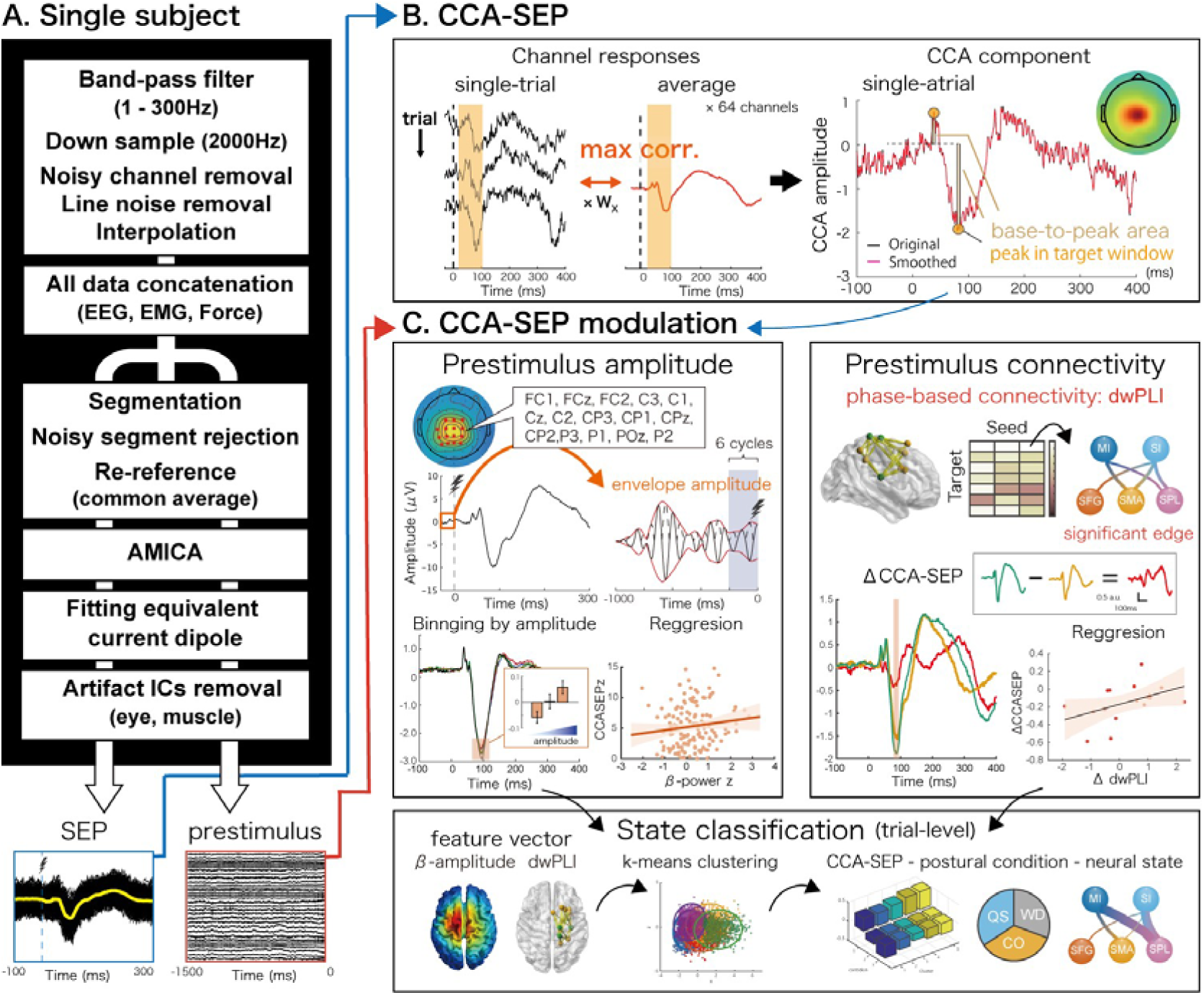
Overview of the analysis workflow and EEG processing pipeline. The left column illustrates the EEG preprocessing for a single participant. The two epochs, the SEP and prestimulus epoch, were finally constructed at the end of these steps. The cleaned EEG data from the SEP epochs were used to compute CCA-SEPs and to quantify the P40 and N80 components for each trial. The prestimulus EEG data were subjected to the Hilbert transform, and the envelope amplitudes at the α-and β-frequency bands were quantified. For connectivity analysis, the time course of source activities was calculated using the source-localisation approach. ΔdwPLI, the difference from the reference SIT condition, was used as the connectivity strength to test the contribution of the cortical connectivity to SEP modulation. In addition, the clustering approach was performed using the relative value of β-amplitude and α-connectivity to those in SIT to examine their association with CCA-SEP modulation.

The continuous EEG time series was segmented into two epochs: the SEP epoch and the prestimulus epoch (Figure 1A). The SEP epoch ranged from −100 to 400 ms relative to the onset of the electrical stimulation, whereas the prestimulus epoch extended from −1,500 to 0 ms. These two epochs were analysed separately in the following steps. The epoch data were applied to automatic rejection (exceeding the average ± 5 standard deviations) and re-referenced to the common average. Subsequently, an adaptive mixture-independent component analysis was applied to the EEG epochs to distinguish between brain and non-brain components for the reduction of the residual artifacts. Finally, a single equivalent current dipole was calculated for the scalp projection of each independent component using a standard MNI model. For each participant, the cleaned EEG time courses from both epochs were standardised into z-scores at each point across all data.

#### Single-trial SEP extraction

For the single-trial SEP analysis, canonical correlation analysis (CCA) was applied to the EEG time courses and their averaged waveforms (Figure 1B, left panel). First, the EEG data were low-pass filtered at 45 Hz using a fourth-order zero-phase filter, following a previous study (Stephani *et al*., 2022). Using matrices *X*, the concatenated EEG of all epochs in the time domain, and *Y*, the average SEP waveform repeated for each epoch, CCA identified the linear transformation weights *Wx* and *Wy* that maximised the correlation between the canonical variates *XWx* and *YWy*. To capture the early SEP components, CCA weights were estimated using data from 30 to 100 ms after stimulation. The CCA analysis yielded a set of weight vectors (*Wx*), each consisting of one weight per EEG channel, with the corresponding components ranked according to their canonical correlation coefficients. The SEP-related CCA component was selected based on both a high canonical correlation coefficient and its spatial activation pattern obtained from covariance-transformed CCA weights (Haufe *et al*., 2014). Specifically, we selected a component whose activation pattern exhibited a central distribution over the foot representations of the sensorimotor cortex and closely resembled the electrode-level SEP topography at the latency of interest. The CCA waveform was obtained by projecting the EEG data matrix onto the selected *Wx*. The polarity of the waveform was adjusted, when necessary, by inverting the sign of the *Wx* so that the P40 and N80 appeared as positive and negative peaks, respectively.

To quantify the SEP component derived from CCA (CCA-SEP), the peak latencies of the earliest response (P40) and the subsequent response (N80) were identified on a single-trial basis (Figure 1B, right panel). For each participant, the search window for these peaks was defined with reference to the peak latencies in the averaged waveform, which was obtained by averaging all signals across all conditions. For each component, a ±3 ms time window centred on the corresponding peak latency in the averaged waveform was used. To reduce high-frequency fluctuations, single-trial waveforms were smoothed using a moving median filter with a 3-ms window. For the baseline period, a reference time point preceding the rising phase of P40 was identified from the average waveform across all conditions. The baseline CCA-SEP was defined as the mean amplitude during the 10-ms preceding this reference point for each trial. This baseline value was then subtracted from the single-trial CCA-SEP waveform. The baseline-corrected response was quantified by integrating the waveform over a 5-ms window centred on the peak latency. The signed area was calculated using the trapezoidal rule and divided by the duration of the time window to obtain the mean response amplitude. The quantified CCA-SEP amplitudes were standardised to z-scores within each participant across conditions. For visualisation, the time courses of CCA-SEPs were also standardised at each time point across all data within each participant. The same procedure was applied to the SEP recorded at Cz (Cz-SEP) as a channel-level reference, and the quantified Cz-SEP amplitudes were also standardised in the same manner.

To evaluate the validity of single-trial analysis, the signal-to-noise ratio (SNR) was calculated for the CCA-SEPs. For each trial, the peak amplitude of the CCA-SEP was divided by the standard deviation of the baseline period of the CCA waveform. The single-trial SNR was then averaged within each condition and the P40 and N80 components were compared. The SNR at Cz was used as a channel-level reference and compared with that of the CCA-SEP. To confirm the spatial distribution of the SEP-related SNR across electrodes, the coefficient of determination (r^2^) was further calculated for each electrode based on the relationship between the single-trial amplitudes and binary labels indicating baseline or SEP peak responses (Cacace & McFarland, 2003).

#### Prestimulus oscillatory amplitude

To characterise prestimulus oscillatory activity, envelope amplitude was computed in the α- and β-frequency bands (Figure 1C, left panel). Because the SEPs exhibited a spatially restricted topography, electrodes used to quantify envelope amplitude were selected based on the activation pattern of the SEP-related canonical variate in electrode space. As described above, the CCA weights were transformed into activation patterns using a covariance matrix. For each participant and condition, electrodes showing high activation with the same polarity as the N80 component were identified, and one point was assigned to each electrode ranked within the top 10% of electrodes. The resulting scores were then aggregated across all participants and conditions using a voting procedure.

The duration of the prestimulus EEG waveform was defined as six cycles of the lowest frequency in each frequency band (that is, alpha: 750 ms for 8 Hz, beta: 462 ms for 13 Hz). The segmented EEG was mirrored on both sides to avoid edge artifacts and filtered with a 4th-order Butterworth filter in forward and backward directions. The analytical signal was then obtained using the Hilbert transform, and its absolute value was taken as the amplitude envelope. The envelopes were averaged and log-transformed in each channel. Finally, the envelope amplitudes across the selected channels were linearly combined using the corresponding CCA weights and standardized to a z-score across all conditions within every participant.

#### Prestimulus corticocortical connectivity

Since SEPs are generated by neural activity in a relatively restricted set of cortical regions (Allison *et al*., 1989; Hamalainen *et al*., 1990; Forss *et al*., 1994; Baumgartner *et al*., 1998; Hari & Forss, 1999; Hu *et al*., 2012), a seed-based connectivity approach was used (Figure 1C, right panel). For the source-level analysis, seed ROIs were defined in the primary motor cortex (M1), primary somatosensory cortex (S1), and secondary somatosensory cortex (SII). Although the networks underlying SEP modulation are not fully understood, the target regions were selected based on their potential involvement in sensory gating and tactile perception within frontal and parietal regions (Kristeva-Feige *et al*., 1996; Grunwald *et al*., 2003; Hauck *et al*., 2006; Boly *et al*., 2007; Vallinoja *et al*., 2021; Hssain-Khalladi *et al*., 2024): the supplementary motor area (SMA), middle frontal gyrus (MFG), inferior frontal gyrus opercular part (IFGop), triangular part (IFGtri), superior frontal gyrus (SFG), superior parietal lobule (SPL), and supramarginal gyrus (SMG). The predefined ROI approach enabled a hypothesis-driven analysis while enabling an exploratory assessment of connectivity patterns across candidate regions and maintaining interpretability and statistical robustness.

To localise source activity for corticocortical connectivity, a standard boundary element method head model was used, and electrode positions were coregistered to the template head. The source space was defined on a standard MNI volumetric grid with a resolution of 10 mm. Source reconstruction was performed using an LCMV beamformer with a covariance matrix computed from the segmented EEG data using 5% regularisation. To define atlas-based ROIs, each grid point was assigned an anatomical label according to the AAL atlas. For each of the 10 seed and target ROIs, the grid point showing the most negative peak within the N80 time window, as determined from the electrode SEP analysis, was identified. A spherical ROI with a radius of 15 mm was defined around this point and constrained within the corresponding anatomical label, thereby anchoring each ROI to putative SEP-related regions. Source-reconstructed time series within each ROI were summarized using principal component analysis (PCA) after polarity correction, and the first component was used as a representative ROI signal for connectivity analysis. In addition, a source map of the SEP response was separately estimated using minimum-norm estimation for visualisation. These procedures were performed using Fieldtrip and BrainNet Viewer implemented in MATLAB (Oostenveld *et al*., 2011; Xia *et al*., 2013).

For these ROI signals, the duration of the prestimulus interval was set to 12 cycles of the lowest frequency in the alpha and beta frequency bands (that is, alpha: 1,500 ms for 8 Hz, beta: 920 ms for 13 Hz). After mirroring the signals at both ends, a fourth-order Butterworth filter was applied in forward and backward directions. Connectivity strength between the seed-target pair was evaluated using the debiased weighted phase lag index (dwPLI; debiased wPLI-square estimator) (Vinck *et al*., 2011):

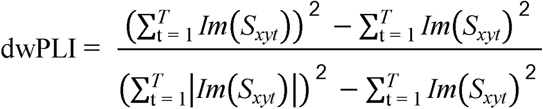

where *T* is the total number of time points, *Im*(*S_xyt_)* denotes the imaginary part of the cross-spectrum at time *t*. Using 3 seed and 7 target regions, dwPLI was calculated for 21 seed-target pairs (edges) in α- and β-frequency bands. To compare across conditions, dwPLI values were standardised to z-scores within each participant before statistical analysis. Repeated-measures correlations were further performed between all pairs of edges to assess shared variation among connectivity features. In addition to examining condition-dependent differences in individual edges, dwPLI values were averaged across all 21 pairs to obtain an edge-averaged global dwPLI, which was used to assess aggregate changes in functional connectivity across predefined connections (Hardmeier *et al*., 2014).

#### Amplitude-related and connectivity-related CCA-SEP modulation

According to previous research (Nikouline *et al*., 2000; Stephani *et al*., 2022), the effects of prestimulus envelope amplitude on lower-limb CCA-SEP were investigated using a linear mixed-effects model (LME). In addition, the trial-level data were binned into four bins within each participant according to prestimulus envelope amplitude, and the lowest (first) and highest (fourth) bins were compared using paired t-tests. In addition to this trial-by-trial analysis, condition-averaged SEPs were analysed to examine whether prestimulus cortical connectivity shaped by the postural demands of each condition, was associated with somatosensory response.

For corticocortical connectivity, we examined whether connectivity patterns modulated by postural demands covaried with SEP modulations. To obtain stable estimates of connectivity strength, dwPLI values were averaged within each condition for each participant. First, edges relevant to task-dependent differences were identified by statistically comparing dwPLI across conditions. We then assessed whether changes in the connectivity strength relative to the reference condition (ΔdwPLI) across these task-sensitive edges were associated with changes in the CCA-SEP (ΔCCA-SEP). Both ΔdwPLI and ΔCCA-SEP were calculated relative to SIT to reduce the influence of edge- and participant-specific differences in absolute baseline values and to emphasise state-dependent modulation associated with postural conditions. As selected edges constituted a high-dimensional feature set and showed substantial inter-edge correlations (Supplement Figure1), PCA was applied to extract the dominant pattern of connectivity variation. Finally, LME was used to test whether the first principal component (PC_1_) was associated with ΔCCA-SEP.

#### Clustering of single-trial epoch with prestimulus oscillatory amplitude and connectivity

To examine whether multivariate patterns of prestimulus neural activity could account for SEP modulation, we performed unsupervised clustering of trial-level feature vectors (Figure 1C, bottom panel). Subsequent analyses examined whether the occurrence probability of each state differed across conditions, whether SEP amplitudes differed across states and conditions within each state. Feature vectors for this classification were constructed from β-amplitude and α-band dwPLI, both of which were significantly associated with CCA-SEP. For each participant, both variables were expressed as differences relative to the SIT condition and standardised across conditions. PCA was applied to the dataset without condition labels, and the number of PCs accounting for more than 60 % of the total variance was retained to define the reduced feature space. K-means++ was then performed in this reduced space with 200 replicates and a maximum of 1,000 iterations to ensure stable clustering solutions. The optimal number of clusters was determined using the elbow method. The CCA-SEP amplitude and proportion of trials for each condition were subsequently evaluated across the resulting clusters. To validate clustering stability, the dataset was repeatedly divided into two halves, stratified by participants and conditions to minimise imbalance between the two subsets (Meilă, 2007). K-means++ clustering was performed on one subset, and the resulting cluster centroids were used to assign observations in the other subset to clusters. The consistency between clustered labels was evaluated over 100 repetitions using the adjusted Rand index.

### Statistical analysis

Predictor variables entered into the LME were decomposed into within- and between-participant components using person-mean centring (Curran & Bauer, 2011). The within-participants component was subsequently standardised to z-scores within each participant, whereas the between-participants component was standardised across all participants. Because the primary focus was on within-participant associations, between-participant effects were not interpreted further. Degrees of freedom for the fixed effects were estimated using the Satterthwaite approximation to derive p-values.

To confirm the validity of single-trial CCA-SEP analysis, SNRs were compared between the CCA-SEP and Cz-SEP methods for both P40 and N80 components. An LME was fitted with method (CCA-SEP vs. Cz-SEP), peak (P40 vs. N80), and their interaction as fixed effects, with random intercepts for participants and participant-by-condition combinations. The significance of the fixed effects was assessed using an analysis of variance (ANOVA).

To test the effect of prestimulus oscillatory amplitude CCA-SEP, LME models were fitted separately for the α- and β-frequency bands. Based on the SNR analysis, which showed a higher SNR for the N80 component, the subsequent analysis was focused on the N80 amplitude of the CCA-SEP. Standardised CCA-SEP amplitude was entered as the dependent variable, and standardised prestimulus amplitude, postural condition, and their interaction were included as fixed effects. A random intercept for participants was included to account for repeated measures. For the two standing conditions, QS and WD, COPv was additionally included as a covariate in the LME to examine whether it was independently associated with CCA-SEP and whether the effects of prestimulus amplitude remained after accounting for COPv. For prestimulus oscillatory amplitude, COPv, and SOL-RMS, condition differences were further tested using an LME model.

To define a feature set of connectivity edges for the analysis of connectivity-related CCA-SEP modulation, condition differences in standardised dwPLI were assessed using an LME for each edge. The resulting p-values for the main effect of the postural task were corrected for multiple comparisons across all seed-target pairs using the false discovery rate (FDR). Edges with FDR-adjusted p-values < 0.1 were retained to broadly capture candidate connections, including weaker but potentially coordinated connectivity changes, for subsequent dimensionality reduction. PCA was then applied to the selected edges. PCs accounting for more than 60.0% of the total variance were retained. In the current dataset, this criterion resulted in the retention of PC1 alone. Subsequently, an LME was fitted with PC1, postural conditions, and their interaction as fixed effects and ΔCCA-SEP as the dependent variable. A random intercept for participants was included to account for repeated measures, which was consistent with the prestimulus amplitude analysis. In addition, global dwPLI and condition differences were also tested using an LME model.

In the trial-level analysis using k-means ++ clustering, mean CCA-SEP amplitude and mean COPv were compared across the resulting clusters (states) using separate LME models. Each model included state as a fixed effect and participant as a random intercept. To determine whether the occurrence probability of each state differed among the four experimental conditions, a separate binomial generalised linear mixed model was fitted for each state. The condition was included as a fixed effect, with a random intercept for each participant. The overall effect of condition was assessed using a Wald χ^2^ test, and pairwise comparisons between conditions were performed on the logit scale using z tests with Tukey adjustment. Estimated marginal means are presented as probabilities on a response scale with asymptotic 95% confidence intervals.

Post hoc pairwise comparisons of SNR, CCA-SEP amplitude, prestimulus amplitude, COPv, SOL-RMS, and global dwPLI were adjusted using the Bonferoni method. For measures in the clustering analysis, p-values from multiple comparisons were adjusted using the FDR procedure. Adjusted p-values are reported throughout the manuscript. The significance level was set at α = 0.05 for all statistical tests. All preprocessing was performed using MATLAB software (2025b), and statistical analyses were conducted using RStudio (ver. 2025.05.1+513).

## Results

### CCA-SEP, signal-noise ratio, and COP velocities across postural tasks

Figure 2A shows the representative averaged SEP waveforms and scalp topography for a single participant. The P40 and N80 components showed maximum amplitudes at Cz, which overlays the cortical representation of the lower limb, as confirmed by topography. For the average CCA-SEP amplitude in N80, one-way ANOVA reveals a significant main effect of postural task (F_3,64_ = 30.226, p < 0.001, partial η^2^ = 0.586). Post hoc comparisons revealed significant differences between all condition pairs except SIT vs. QS, and CO vs. WD (all significant pairwise comparisons, Bonferroni-adjusted p < 0.001, Figure 2C, N80). CCA-SEP was significantly decreased in the CO and WD conditions compared with the other postural conditions. Similar results were obtained for Cz-SEP (Supplement Figure2).

**Figure 2.**
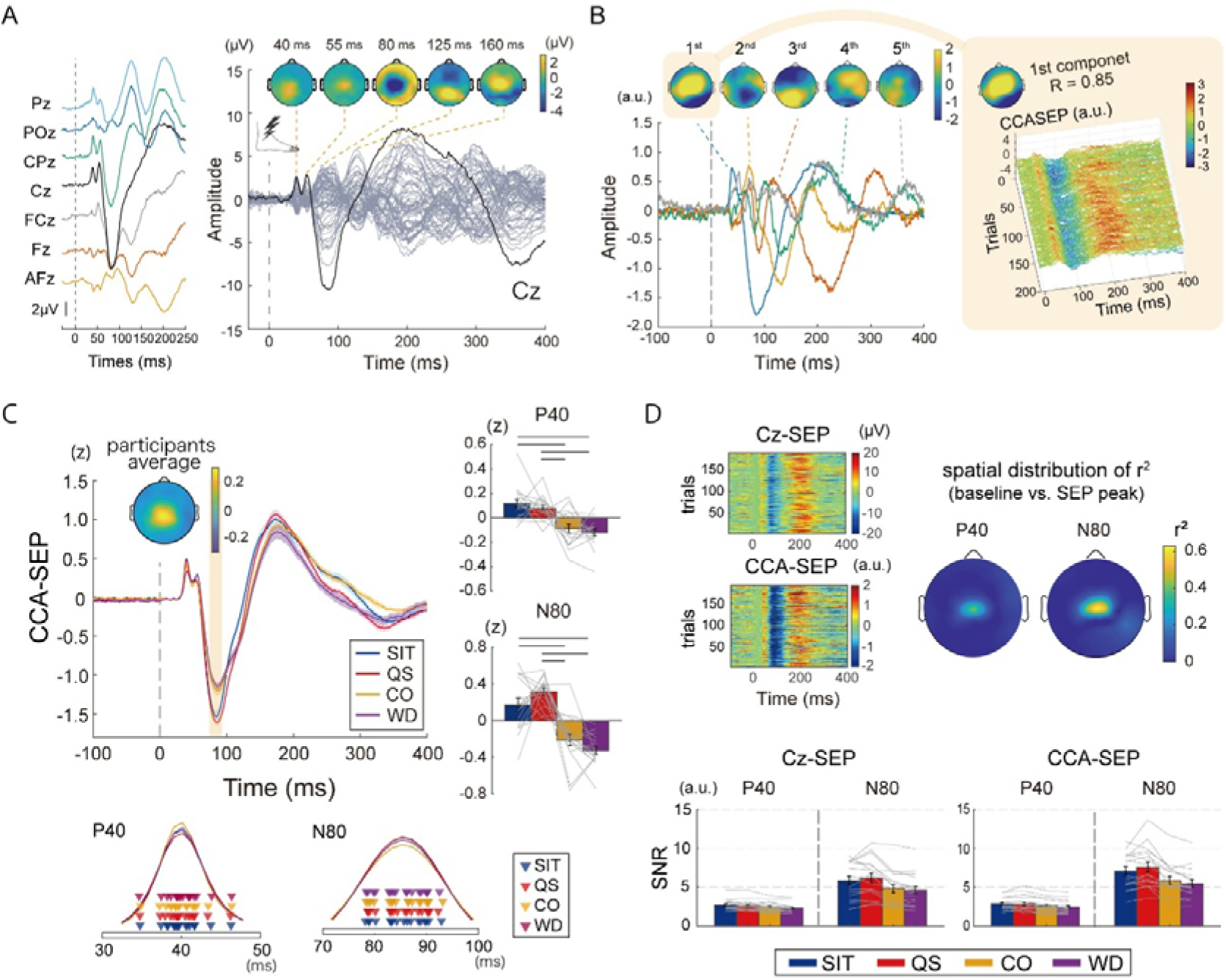
Channel-level SEP and CCA-SEP in different postural tasks. A. Example time course of the averaged SEP at representative midline electrodes and butterfly plot for a single participant in the SIT condition. Topographic maps of scalp potentials at the indicated time points are shown on the butterfly plot. The Cz waveform is highlighted as a typical lower-limb SEP. B. CCA components of a representative participant in the QS condition. Five components, ranked by their coefficients within the target SEP window, are shown. In this case, the first component was selected because it showed a high correlation coefficient and a centrally distributed large amplitude, consistent with the SEP at around 80ms. Its stability is also visually apparent in the trial-by-trial waveforms. C. Averaged SEP waveform in all participants for the CCA-SEP. Quantified P40 and N80 are statistically compared across four conditions. Peak timings of both components are depicted for each participant. No significant difference was observed in the peak timing across conditions. D. SNR analysis for the SEP-CCA and Cz-SEP. *upper*; trial-by-trial SEP on the time-amplitude space in the QS condition from a representative participant. The spatial distribution of the coefficient of determination (r^2^) is also presented at the timing of P40 and N80 in the same participant. Consistent with the SEP topography, the central electrodes showed higher r^2^, indicating better SNR. *lower*; averaged SNR for peak P40 and N80 relative to the standard deviation of baseline amplitude in all participants.

Similarly, in the earlier component (P40) of the average CCA-SEP, a significant main effect of the postural task (F_3,64_ = 14.23, p < 0.001, partial η^2^ = 0.400) was observed, and post hoc comparisons revealed significant differences between the SIT and the CO, SIT and WD, and the QS and the CO, and the QS and WD. As with N80, the earlier component of CCA-SEP was significantly decreased in the CO and WD conditions compared with SIT and QS conditions (Figure 2C, P40).

A significant interaction between method and peak was observed for SNR (F_1,201_ = 10.75, p = 0.00123, partial η^2^ = 0.0507). Post hoc analysis revealed that the SNR was higher for CCA-SEPs than for Cz-SEPs at N80 (p < 0.001), whereas no differences were observed at P40 (p = 0.340). In both methods, the SNR was significantly higher at N80 than at P40 (both p < 0.001), indicating that the N80 provides a more robust component for single-trial analysis. Since N80 exhibited greater SNR (Figure 2D) and effect size of peak (p < 0.001, partial η^2^ = 0.725), subsequent analysis focused on the N80 component in the CCA-SEP.

For the COPv compared between QS and WD conditions, the main effect of postural task was significant (F_1,16_ = 10.62, p = 0.00492, partial η^2^ = 0.399), showing that averaged COPv was significantly higher in the WD condition. For the SOL-RMS, a main effect was observed in postural task (F_3,64_ = 48.77, p < 0.001, partial η^2^ = 0.695). Post hoc comparisons revealed significantly lower activity in the SIT condition than in the other conditions (all p < 0.001) and significantly higher activity in the WD condition than in the SIT (p < 0.001), QS (p = 0.001), and CO (p = 0.008) conditions. Notably, no significant difference was observed between the QS and CO conditions (p = 1.00), indicating that the contraction task in the CO condition was performed as intended.

### Prestimulus amplitude-related CCA-SEP modulation

To calculate prestimulus amplitude, 15 electrodes over the central area with the highest voting scores were selected (Figure 3A). In the β-frequency band, two-way ANOVA using the LME revealed a significant interaction between prestimulus β-amplitude and postural task for within-participants (F_3, 10212_ = 3.21, p = 0.0217). Simple slope analyses showed that the slopes were significant in the QS, CO, and WD conditions, but not in the SIT condition (Bonferroni-adjusted p = 0.706; QS: b = 0.0477, Bonferroni-adjusted p = 0.0110; CO: b = 0.0432, Bonferroni-adjusted p = 0.0218; WD: b = 0.0760, Bonferroni-adjusted p < 0.001). The variance of the random intercept was estimated to be near zero, and the model exhibited a singular fit. Therefore, as a sensitivity test, we applied a linear model with participant-clustered robust standard errors. The results showed the interaction was still significant between the prestimulus β-amplitude and postural task (F_3, 10212_ = 5.05, p = 0.00169), and the slope in the SIT condition was close to zero (b = −0.00712), whereas the slope in the WD condition was positive (b = 0.0760), and the difference between these slopes was significant (Bonferroni-adjusted p = 0.0149). Collectively, these findings support the robustness of the main analysis with the LME. In the bin-based analysis, CCA-SEP amplitude was significantly larger in the fourth bin, which had the highest mean prestimulus β-amplitude, than in the first bin, which had the lowest mean prestimulus β-amplitude (t(16) = 4.97, p < 0.001). Figure 3B shows the association between prestimulus β-amplitude and CCA-SEP and shows single-trial waveforms from a representative participant. In the two standing conditions (QS and WD), prestimulus β-amplitude remained positively associated with CCA-SEP amplitude after accounting for postural sway (b = 0.050, F_1, 4897_ = 13.98, p < 0.001, Supplement data), whereas COPv showed a negative association (b = −0.032, F_1, 4897_ = 5.03, p = 0.0248).

**Figure 3.**
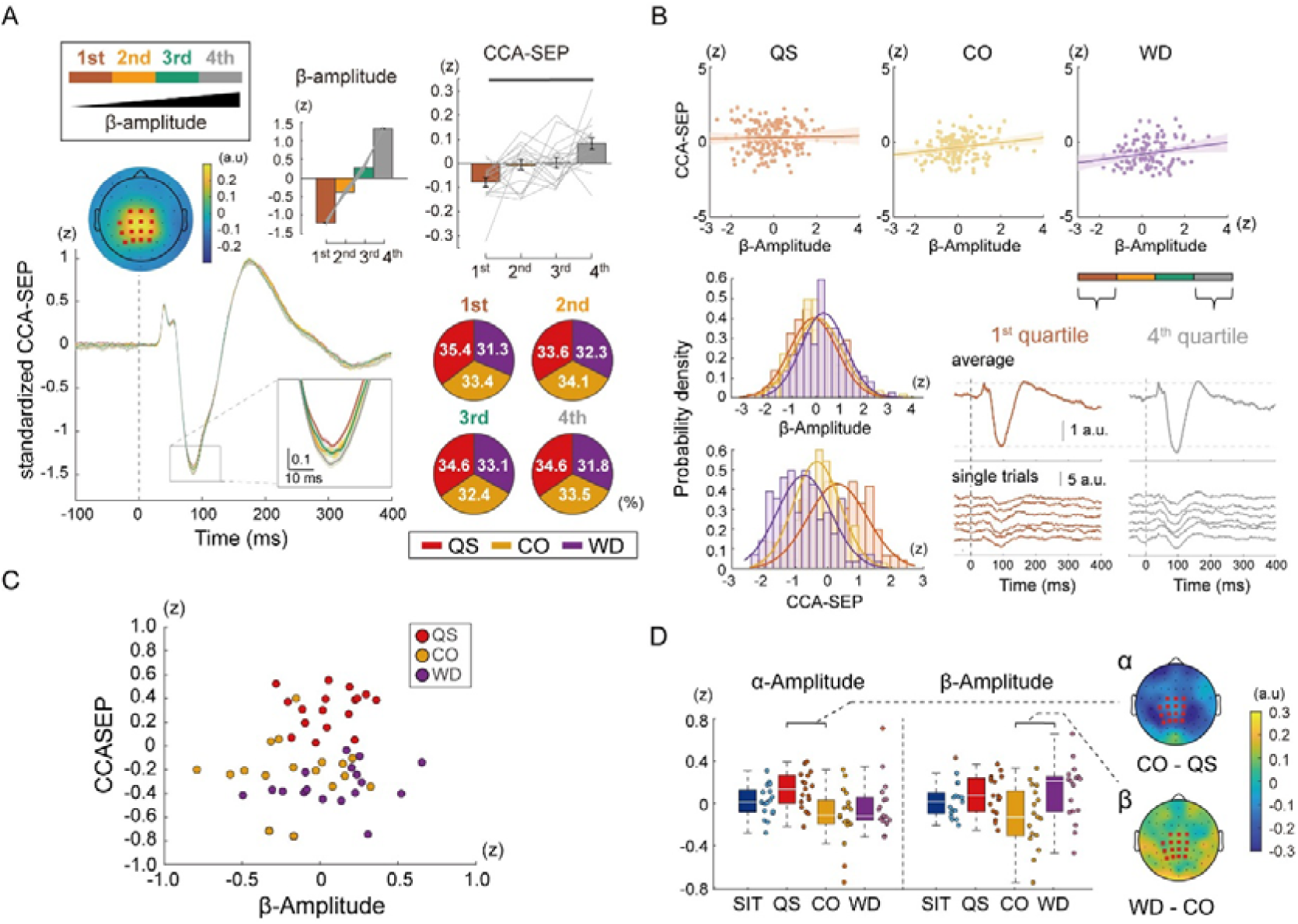
Amplitude-related modulation of lower limb CCA-SEP. A. CCA-SEP in four subgroups binned according to prestimulus β-amplitude. Binning was performed separately for each participant. The bar graph indicates a positive relationship between β-amplitude and CCA-SEP amplitude: the first bin, corresponding to low β-amplitude, shows a smaller CCA-SEP, whereas the fourth bin, corresponding to high β-amplitude, shows a larger CCA-SEP. For visualisation, CCA-SEP waveforms were pooled across the QS, CO, and WD conditions and averaged across all participants. The 15 electrodes used for prestimulus amplitude analysis are highlighted on the averaged CCA-SEP topographic map. In addition, the pie charts show the proportions of the four conditions within each bin across all participants. B. Significant association between the CCA-SEP and prestimulus β-amplitude in the QS, CO, and WD conditions. Upper panels show the association between CCA-SEP and prestimulus β-amplitude in representative participants, with positive regression slopes observed in all three conditions. In the WD condition, trials were binned according to β-amplitude. Averaged CCA-SEP waveforms and representative single-trial waveforms are shown for the first and fourth bins. The histograms of CCA-SEP and β-amplitude indicate comparable variabilities across three conditions. C. Relation between condition-averaged prestimulus β-amplitude and CCA-SEP. Each point represents the mean value for each participant and condition. Averaged β-amplitude was not significantly associated with CCA-SEP, indicating that task-dependent differences in the β-amplitude were not significantly associated with averaged CCA-SEP. D. Condition-averaged prestimulus envelop amplitude in the α- and β-frequency bands. Grand-average scalp topographies of the significant between-condition differences are also presented.

For the α-frequency band, the LME model showed a significant main effect of prestimulus α-amplitude (F_1, 10212_ = 5.44, p = 0.0200). However, no significant differences were observed in the simple slopes for each condition. Furthermore, in the sensitivity analysis using the linear model, the effect of prestimulus amplitude did not reach significance (b = 0.0225, t_15.71_ = 2.083, p = 0.0539), indicating the marginal association between prestimulus α-amplitude and CCA-SEP.

To examine the association between the prestimulus oscillatory amplitude shaped by postural demands and the CCA-SEP, an LME was applied to the condition-averaged SEPs and β-amplitude in the three conditions showing significant trial-level associations (QS, CO, WD). Neither interaction (p = 0.332) nor the main effect (p = 0.570) was significant (Figure 3C), indicating that condition-averaged β-amplitude did not clearly account for the variability in condition-averaged CCA-SEP.

Furthermore, we compared the average prestimulus amplitude at the α- and β-frequency bands across the four conditions. For the prestimulus envelope β-amplitude, the main effects were observed on the postural task (F_3,64_ = 3.53, p = 0.0198, partial η^2^ = 0.142), and the post hoc analysis showed a significant decrease in the CO conditions than the WD condition (Bonferroni-adjusted p = 0.0239, Figure 3D). At the α-frequency band, a main effect (F_3,64_ = 3.69, p = 0.0162, partial η^2^ = 0.147) and a significant difference between the QS and CO conditions were noted (Bonferroni-adjusted p = 0.0116).

### Corticocortical connectivity-related CCA-SEP modulation

To examine task-dependent modulation at the level of cortico-cortical connectivity, we quantified connectivity strength using dwPLI and tested whether prestimulus connectivity was associated with SEP modulation across postural tasks. For the edge-averaged global dwPLI, main effects were noted among conditions (F_3, 64_ = 6.18, p < 0.001, partial η^2^ = 0.225), and the post-hoc analysis revealed significant differences between SIT and WD (Bonferroni-adjusted p < 0.001), QS and WD (Bonferroni-adjusted p = 0.0294), and CO and WD (Bonferroni-adjusted p = 0.0266) conditions (Figure 4B). Four significant edges were observed within the 21 seed-target pairs at the α-frequency band (M1-SMA, F_3, 48_ = 4.74, FDR-adjusted p = 0.0294, partial η^2^ = 0.229; M1-SPL, F_3, 48_ = 5.26, FDR-adjusted p = 0.0294, partial η^2^ = 0.248; S1-SMA; F_3, 48_ = 7.85, FDR-adjusted p = 0.00484, partial η^2^ = 0.329; S1-SPL, F_3, 48_ = 4.88, FDR-adjusted p = 0.0294, partial η^2^ = 0.234, Figure 4C). At the β-frequency, no significant main effect was observed in any pair across conditions. Therefore, the following analysis was performed on the α-connectivity.

**Figure 4.**
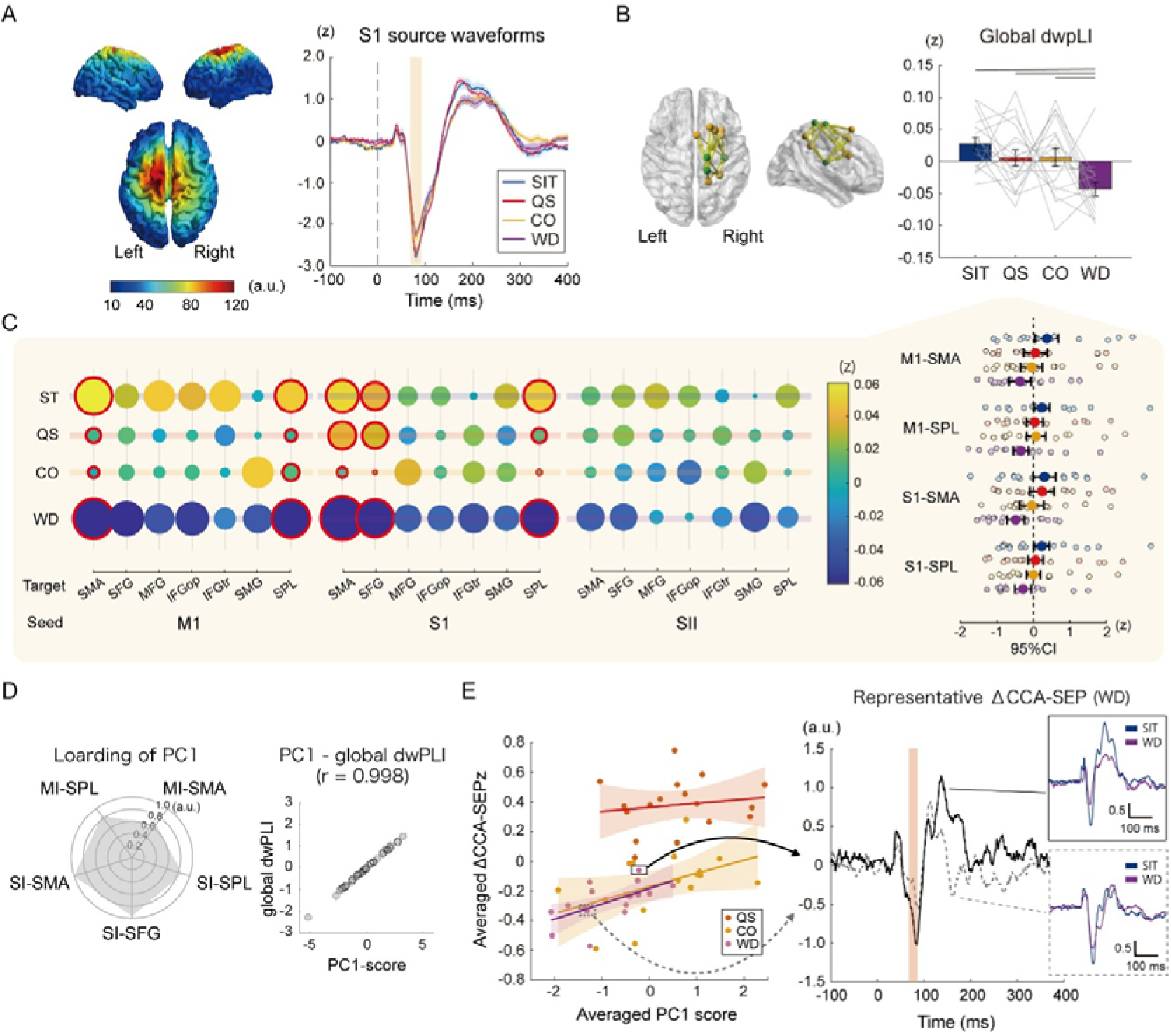
Connectivity-related modulation of CCA-SEP. A. Representative N80 cortical maps in the SIT condition and reconstructed waveforms of each condition in the SI. B. Ten regions of interest for the connectivity analysis and averaged global dwPLI of four conditions in all participants. *green*, three seed regions; *orange*, seven target regions C. Bubble plots for standardised dwPLI. Relative differences in the circle size indicate the absolute value of the dwPLI. Note that the mean value 0 is the smallest. Red circles represent significant differences. CI plots are presented for the significant edges in all participants, replicating the result of the global dwPLI (that is the connectivity strength decreases in the WD condition). *CI*, confidence interval. D. *left*: Radar plot of five edges for PC_1_, which is a representative connectivity strength vector. *right*: Correlation analysis between global dwPLI and PC_1_ scores. Higher PC1 scores were associated with higher mean dwPLI. E. Scatter plot for ΔCCA-SEP and ΔPC_1_ in three conditions. The right panel shows the typical waveform of large and small ΔCCA-SEP in the WD condition.

PCA for identifying the feature vector of cortical connectivity extracted the PC_1_ which explained 61.1% of the variance. The PC_1_ score, which highly correlated with the global wPLI (correlation coefficient = 0.998, Figure 4D), was substituted into the LME as a predictor to test the influence of connectivity strength and condition on the CCA-SEP. Results showed a main effect of changes in connectivity strength with no interaction (F_1,42_ = 6.27, p = 0.0162, partial η^2^ = 0.130), which indicates the positive relation between the modulation of the CCA-SEP and connectivity strength (Figure 4D). No significant difference was noted in the interaction, indicating that this association did not differ across the three conditions (F_2,42_ = 0.631, p = 0.537). These results revealed that CCA-SEP modulation covaried with posture-related changes in α-connectivity relative to the resting seated condition.

### CCA-SEP modulation associated with prestimulus neural states

To examine whether multivariate patterns of prestimulus cortical activity accounted for trial-to-trial SEP modulation, we used an unsupervised clustering approach. The first three PCs derived from the connectivity strength matrix explained above 60% of the total variance (PC_1_, 31.3%; PC_2_, 17.7%; PC_3_, 16.1%). The PC scores and Δβ-amplitude were used as feature vectors for k-means++ clustering. As a result, 7,602 epochs were classified into six clusters based on the elbow points (Figure 5A).

**Figure 5.**
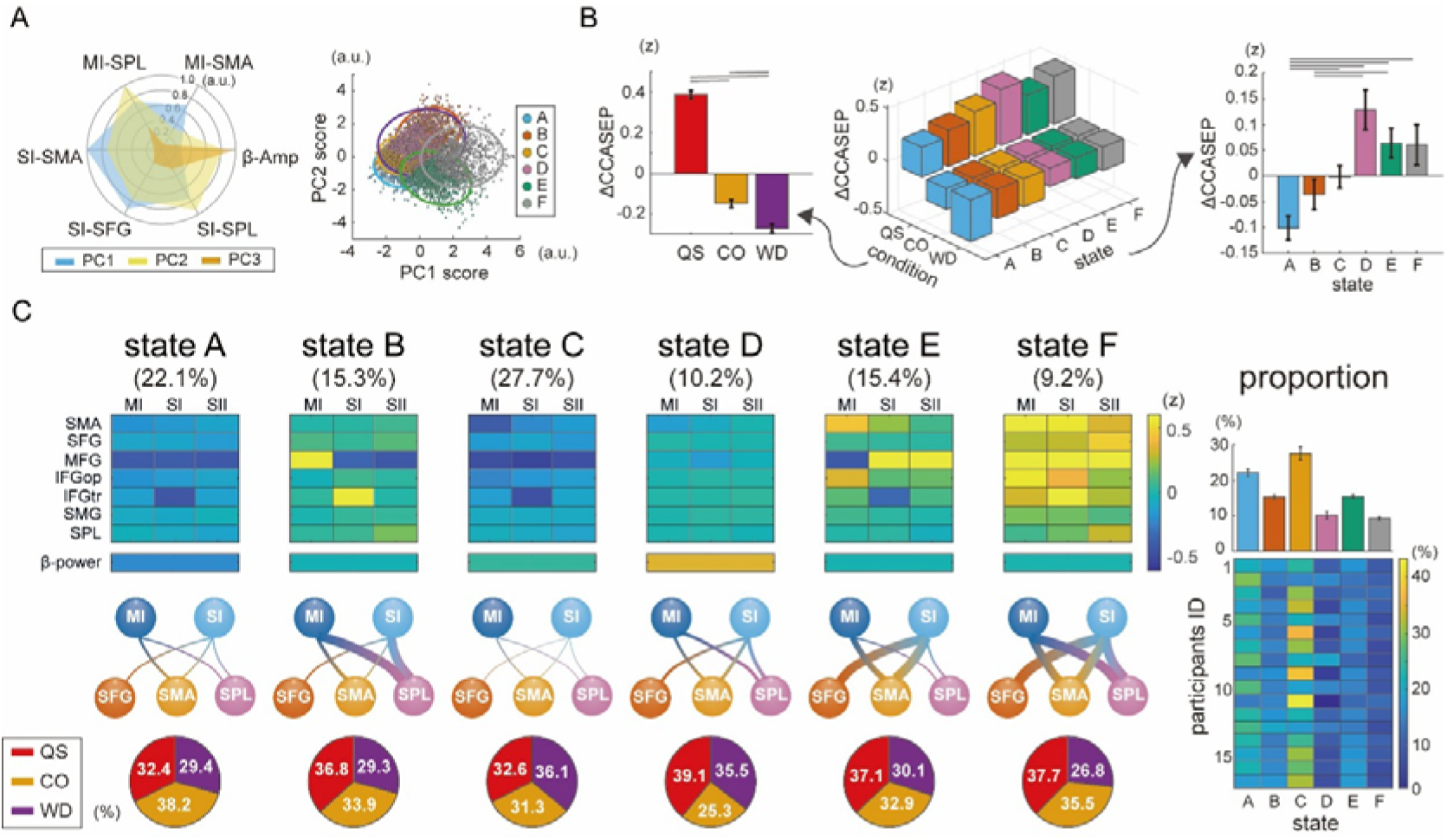
Classification of the neural state with changes in β-amplitude and connectivity. A. *Left*: Radar plot showing the three PC features used for clustering. *Right*: Scatter plot for PC_1_ and PC_2_ scores of the five clusters in two-dimensional space. B. 3D and 2D bar graphs showed the relation among the ΔCCA-SEP, experimental condition, and neural state allocated with clustering. 3D, three-dimensional; 2D, two-dimensional C. Characteristics of feature matrices for the clustering and state probabilities. Matrices indicate the standardised connectivity strength across all 21 pairs. Based on these connectivity strengths, the interregional relationships are schematically presented in the lower panel. Line width indicates the relative ratio between the connectivity strength. Pie charts presenting the proportions of the QS, CO, and WD conditions in each state. The proportion of each state for each participant is shown in the right panel. This confirms that the composition of each state was not dominated by any specific participant.

For the ΔCCA-SEP, statistical tests revealed main effects of the state (F_5,7584_ = 6.74, p < 0.001, partial η^2^ = 0.00442) and those of the postural task (F_5,7584_ = 304.35, p < 0.001, partial η^2^ = 0.0742). ΔCCA-SEP was significantly reduced in state A, followed by state B, in post hoc comparisons (A vs. C, FDR-adjusted p = 0.00200, A vs. D, p < 0.001, A vs. E, FDR-adjusted p < 0.001, A vs. F, FDR-adjusted p = 0.00560, B vs. D, FDR-adjusted p = 0.00200, B vs. E, FDR-adjusted p = 0.0302, Figure 5 B). In the COPv, neither the main effect of state (F_5,176_ = 0.228, p = 0.949) nor the interaction between state and conditions (F_5,176_ = 0.144, p = 0.981) was significant.

Figure 5 C shows the characteristics of each state. State A was characterized by low Δβ-amplitude and weaker connectivity, whereas state D showed high Δβ-amplitude and similar connectivity strength. States B, C, E, and F showed different Δconnectivity with similar Δβ-amplitude. State C exhibited weaker connectivity across all pairs and a higher proportion of WD conditions, whereas state F exhibited greater connectivity and a lower proportion of WD conditions. States B and E showed mirror-symmetric connectivity characteristics, indicating that greater connections were observed in pairs involving the SPL in state B and in pairs involving the SMA and SFG in state E.

Split-half analysis showed high clustering stability (mean ARI = 0.853, SD = 0.085, median = 0.885), indicating that the identified structure was reproducible across repeated random splits. In the proportion of postural tasks in each state, the CO condition occupied a higher proportion in state A and a lower in state D (state A: Wald χ^2^(2) = 24.50, p < 0.001, state D: Wald χ^2^(2) = 24.054, p < 0.001), whereas the WD condition occupied a higher proportion in state C and a lower proportion in state F (state C: Wald χ^2^(2) = 29.35, p < 0.001, state F: Wald χ^2^(2) = 8.568, p = 0.0206).

## Discussion

In this study, we investigated whether transient cortical activity modulates somatosensory processing during postural tasks. The results showed that the N80 component of the lower limb CCA-SEP was significantly reduced when postural demands increased. In addition, a single-trial analysis revealed that prestimulus β-amplitude over the central regions covaries with the N80 in the QS, CO, and WD conditions: Lower prestimulus β-amplitude is linked to a larger reduction of SEP amplitude, and this effect is retained irrespective of the postural sway (COPv). Because the significant relation between β-amplitude and CCA-SEP was not observed within-participant averaged data, this association appears to be driven by moment-to-moment fluctuations in cortical oscillatory activity. Furthermore, we further showed that condition-level modulation of cortical connectivity in the α-frequency band was associated with the CCA-SEP amplitude. Greater reductions in α-connectivity between sensorimotor and frontoparietal regions were associated with stronger somatosensory gating. Finally, the cluster analysis using single-trial multivariate cortical features showed that CCA-SEP amplitude varies across different cortical states characterised by β-amplitude and α-connectivity. These results indicate that responsiveness to somatosensory stimuli is shaped by somatosensory gating associated with postural demands as well as by transient states of ongoing cortical activity.

### Somatosensory gating during postural control

The present results showed that a reduction in the lower-limb SEP is a common feature of both postural control and limb movement and becomes more pronounced with increasing task demands. This is consistent with several studies that reported somatosensory attenuation during postural tasks (Dietz *et al*., 1985; Applegate *et al*., 1988; Restuccia *et al*., 2008). The earliest component, P40, is reduced from a lying posture to normal standing, whereas results are controversial when moving from a sitting position (Dietz *et al*., 1985; Applegate *et al*., 1988). It has been proposed that weight-bearing on the foot is one of the factors responsible for this sensory attenuation in the standing posture (Lhomond *et al*., 2016). In this study, somatosensory attenuation at N80 was enhanced in unstable standing compared with both normal standing and sitting postures, whereas no difference was observed between normal standing and sitting postures. This highlights the functional relevance of somatosensory gating, which targets the unnecessary information to achieve the task goal (that is, stable posture in this study) (McIlroy *et al*., 2003; Wasaka & Kida, 2025). We suspect that the differences in this attenuation across postural tasks partially arise from changes in background neural activity. When posture becomes unstable, cortical excitability is enhanced to execute postural compensation; and spinal reflexes are suppressed (Obata *et al*., 2009). As somatosensory gating is mediated by peripheral mechanisms (Chapman *et al*., 1987; Williams & Chapman, 2002), as well as by top-down control originating from the higher nervous system (Voss *et al*., 2006; Seki & Fetz, 2012), it seems reasonable to propose that differences in background neural activity contribute to variability in the strength of somatosensory attenuation. As potential indices of such background neural activity, associations of CCA-SEP amplitude with the prestimulus β-amplitude and α-connectivity were observed.

Although N80 amplitude did not differ significantly between the CO and WD conditions, the underlying mechanisms may not be the same. The tonic muscle contraction task requires maintaining consistent motor output without large fluctuations, which could dominate the feed-forward control, whereas standing posture is automatically achieved with dynamic reweighting of sensory modality in feed-back control. As observed in the prestimulus β-amplitude and α-cortical connectivity, background neural activity is significantly different between the two tasks. Because somatosensory gating is considered to reflect the net effect of both attenuated and enhanced sensory processing at multiple neural sites, task specificity may not always be evident in reduced SEP amplitudes, which represent the final output of these processes.

Notably, no significant difference in N80 amplitude was observed between the SIT and QS conditions, consistent with classic literature (Dietz *et al*., 1985). As shown in previous studies (Cohen & Starr, 1987; Lin *et al*., 2000) and observed under the CO condition, muscle contraction is a potential factor in somatosensory attenuation. However, although the standing posture required more muscle contraction than the sitting posture, a significant reduction in SEP amplitude was not obvious. This reveals that somatosensory gating cannot be explained solely by the amount of muscle activation but instead reflects an active process through which top-down control modulates sensory integration to support standing balance.

### Trial-to-trial SEP variability associated with prestimulus β-amplitude

The lower-limb CCA-SEP was reduced in the case of the lower prestimulus β-amplitude in the QS, CO, and WD conditions. This relationship was not observed in the sitting posture, indicating a task-dependent effects. Consistent with previous studies in which lower prestimulus α- and β-amplitudes are related to clear modulation of somatosensory responses or tactile perception (van Ede *et al*., 2010; Lange *et al*., 2012; Baumgarten *et al*., 2016; Stephani *et al*., 2020), the present results indicate that somatosensory attenuation is facilitated by the lower amplitude in the β-frequency band. This finding suggests that somatosensory gating fluctuates with ongoing cortical activity, while maintaining posture. Prestimulus oscillations corresponding to the cortical state at the time of sensory stimulation may modulate evoked-potential amplitude through gain control by altering the responsiveness of cortical populations to incoming sensory signals (Becker *et al*., 2008; Iemi *et al*., 2019). In this framework, fluctuations in prestimulus oscillatory activity may change the magnitude of the cortical response elicited by the same sensory input. This interpretation is particularly relevant to early evoked-potential components, which have been proposed to represent additive responses elicited by sensory stimulation, instead of responses generated primarily by the phase resetting of ongoing EEG activity. From this perspective, it is conceivable that SEP amplitude during postural tasks reflects moment-to-moment adjustments of cortical gain as a function of prestimulus cortical activity, in addition to task-dependent attenuation related to postural demands.

Many previous studies highlight the importance of the α-frequency band as a marker of cortical excitability to external events (Baumgartner *et al*., 1998; van Dijk *et al*., 2008; Forschack *et al*., 2017; Iemi *et al*., 2019; Stephani *et al*., 2020). However, in the present study, an association between the prestimulus amplitude and SEP amplitude was observed only in the β-frequency band. Several factors may account for this specific association. First, it may reflect a region-specific frequency band for responsiveness to sensory stimuli. Samaha et al. proposed that α-oscillations are predominant in the visual cortex and relate to the visual stimulus, whereas the β-oscillations are more specific in the parietal cortex, where the somatosensory information is processed (Samaha *et al*., 2017). Thus, the excitability of different cortical regions can be reflected in distinct frequency bands, and the β-frequency band appears to be a gatekeeper in the parietal cortex. Second, the oscillatory activity in the upper α- and lower-β frequency bands may reflect a common aspect of cortical activity. Power in the upper α and neighbouring lower-β bands has been reported to be negatively correlated with broadband high-frequency cortical activity, which is considered an index of local cortical activation or excitability and is closely related to neuronal firing (Manning *et al*., 2009; Iemi *et al*., 2022). Thus, associations with SEP amplitude may emerge within the range spanning the upper α- to lower-β frequency bands.

Third, this may be because β-oscillation has traditionally been interpreted as supporting functional inhibition and the maintenance of the current sensorimotor state (Khanna & Carmena, 2015). More recently, β-oscillations have been proposed to play a role in regulating the representation and updating of internal states or endogenous content (Spitzer & Haegens, 2017; Lundqvist *et al*., 2024). A state in which β-bursts are more likely to occur is considered to suppress information flow within the corresponding regions, thereby transiently maintaining the current internal state. Consistent with this view, near-threshold sensory inputs are less likely to reach conscious awareness when prestimulus β-bursts are prominent (Shin *et al*., 2017; Karvat *et al*., 2021). If higher prestimulus β-amplitude in the present study reflects a greater occurrence of such β-bursts, it may indicate a momentary neural state that is relatively stable and resistant to updating. Such a state may constrain the task-dependent changes in somatosensory processing during postural control, resulting in less somatosensory attenuation and, consequently, a larger SEP amplitude.

### SEP modulation associated with rest-relative changes in α-connectivity

Prestimulus amplitude, power, and cortical connectivity are related to conscious perception of sensory stimuli (Weisz *et al*., 2014; Sadaghiani *et al*., 2015). The present results showed that connectivity strength was significantly weakened in the WD condition, particularly for edges S1-SMA, M1-SMA, S1-SPL, and M1-SPL. In an unstable standing posture, it may be advantageous to enable flexible reconfiguration of networks involving subcortical and cerebellar regions instead of maintaining a fixed, tightly coupled state. Since previous evidence indicates that reduced or altered functional connectivity can be associated with reduced sensory perception and attenuated sensory responses (Weisz *et al*., 2014; Sacchet *et al*., 2015; Sadaghiani *et al*., 2015), the reduced connectivity observed in the WD condition, in which somatosensory gating was enhanced, represents a network state that favours somatosensory gating.

Furthermore, the magnitude of connectivity differences was significantly correlated with SEP modulation in the QS, CO, and WD conditions. Given the positive correlation between PC score and global dwPLI, this result suggests that greater reductions in connectivity strength within individuals are associated with stronger somatosensory gating. Importantly, this connectivity-related modulation was observed only in the α-frequency band, but not in the β-band. Alpha-band connectivity contributes to large-scale connectivity among distinct cortical regions, which mediates motor control as well as cognitive functions such as selective attention, prediction, memory, and executive control (Sadaghiani & Kleinschmidt, 2016). Attention to task-relevant sensory inputs is a well-known factor that facilitates SEP responses (Staines *et al*., 2002; Schubert *et al*., 2008). In the CO condition, participants were required to adjust the contraction level, which may have shifted their attentional resources toward task achievement and away from electrical stimulation. As a result, attentional effects may contribute to the reduction in SEP amplitude as much as in the WD condition. In addition, α-connectivity has been proposed to play an active role in top-down control, particularly in regulating the timing of information processing (Pascucci *et al*., 2025). Alpha-band connectivity contributes to the organisation of rhythmic cycles of cortical excitation and inhibition, thereby determining when sensory inputs are likely to be processed. The present findings are consistent with this view, indicating that reduced α-connectivity reflects a redistribution of sensory-processing networks in response to postural demands. Accordingly, the reduction in SEP amplitude may reflect a simple adjustment of somatosensory gain as well as differences in higher-order cognitive states associated with postural demands.

In the current analysis, we used an exploratory approach in which cortical regions potentially involved in SEP modulation were predefined and connectivity changes among these regions were summarised using PCA. Although this approach does not identify the contribution of a specific connection, it enables us to capture the shared variance across correlated connectivity measures and examine whether this common pattern is associated with SEP modulation. The connectivity identified in this study is unlikely to be the single factor that fully explains differences in SEP reduction across conditions. Because somatosensory gating also occurs at the spinal and subcortical levels (Insola *et al*., 2004; Seki & Fetz, 2012), the final magnitude of gating is likely to be determined by multiple contributing mechanisms. In addition, later SEP components originate from a wider range of cortical regions and are potentially influenced more by cognitive neural networks, indicating that the association between prestimulus connectivity and SEP modulation varies across SEP components.

### SEP variability across prestimulus cortical states

In the clustering analysis applied to 7,602 single-trial epochs over participants and conditions, the states A and B showed significantly smaller SEP responses, further supporting the relevance of prestimulus β-amplitude and α-connectivity in SEP modulation. A comparison between states B and E was particularly informative, because these states showed similar changes in β-amplitude but contrasting changes in α-connectivity between M1/S1 and SFG, SMA, and SPL. This finding highlights again that trial-level SEP modulation is influenced by β-amplitude as well as by α-connectivity. Focusing on the CO and WD conditions, both of which showed reduced SEP amplitudes, the CO condition was characterised by a higher occurrence of state A, which exhibited the lowest β-amplitude, whereas the WD condition was characterised by a higher occurrence of state C, which exhibited weaker connectivity among cortical regions. These findings highlight the differences in the predominant states between the two conditions and support the notion that SEP reduction under the CO and WD conditions may be mediated, at least in part, by different mechanisms. In addition, although the occurrence probabilities of the six clusters were biased across experimental conditions, all conditions were observed within each cluster. Because SEPs were evoked randomly during each condition, this finding implies that cortical states are time-varying during postural control, and responsiveness to somatosensory inputs varies with those transient activities.

Even after cluster classification, a significant main effect of the postural task remained, as shown in Figure 5, indicating that SEP amplitude differed across conditions. This suggests that the neural states identified based on the multivariate cortical features examined in the present analysis are not sufficient to fully explain posture-related somatosensory gating. Instead, they captured trial-to-trial variability in SEP amplitude. Furthermore, no significant difference was observed in COPv among the clusters, indicating that these neural states could not account for differences in postural sway. Revealing novel factors involved in SEP modulation and their functional relevance is an important objective for future research.

## Conclusion

The present study investigated whether somatosensory attenuation during postural tasks reflects posture-related gating as well as fluctuations in cortical activity. Results showed that SEP was reduced more pronouncedly under higher postural demands and that, in single-trial analysis, SEP amplitude positively correlated with prestimulus β-amplitude. Prestimulus α-connectivity, involving sensorimotor regions, also covaried with SEP modulation relative to the seated condition, with relatively greater connectivity associated with more pronounced somatosensory gating. These findings reveal that somatosensory processing for postural control reflects both posture-related gating and moment-to-moment fluctuations in ongoing cortical states. Clarifying how these time-varying changes are related to postural stability may improve our understanding of the neurophysiological mechanisms underlying stable postural control.

## Supplementary materials

### Correlation matrix of 21 edges of cortico-cortical connectivity

**Supplementary Figure 4.**
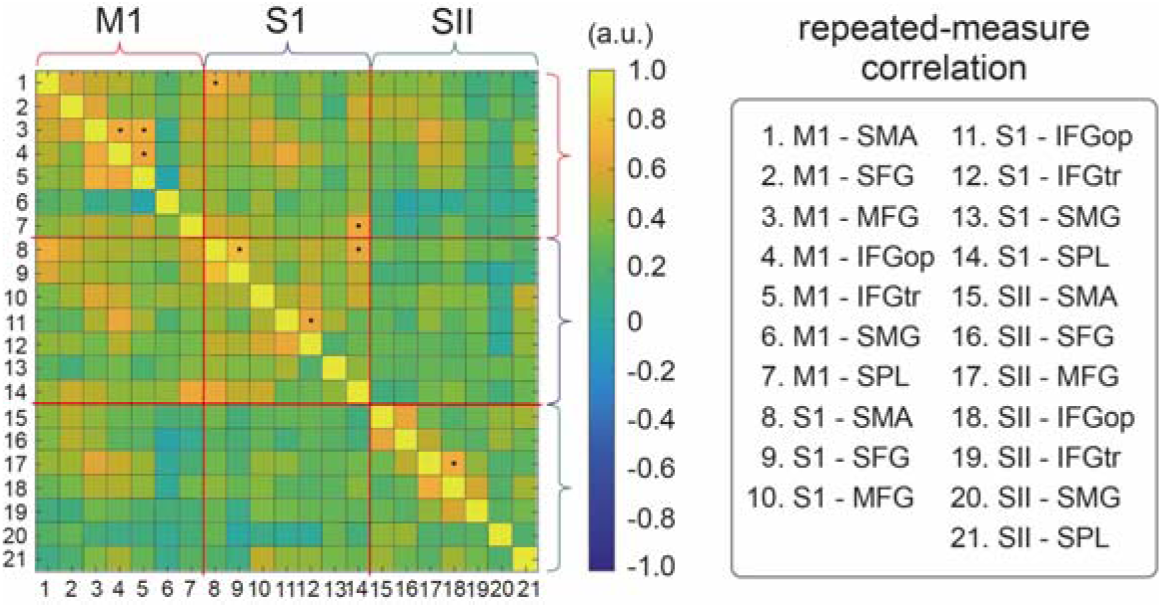
Inter-edge correlation matrix. Before PCA of cortico-cortical connectivity in the α-band, we tested whether connectivity strength showed significant correlations across all pairs. Black dots indicate significant edge correlations based on repeated-measures correlation analysis with Benjamini-Hochberg FDR correction. The threshold for a correlation coefficient was set at 0.65.

### Cz-SEP

**Supplementary Figure 1.**
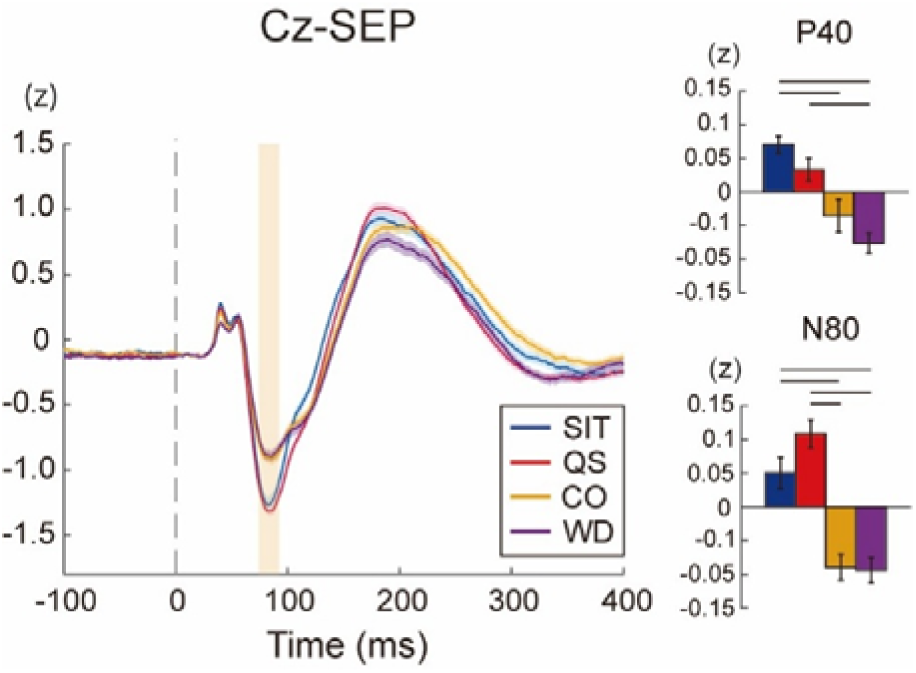
Cz-SEP in different postural conditions. To confirm similarity between the CCA-SEP and SEP at the Cz electrode, a representative electrode for lower-limb SEP, the condition-dependent modulation of the Cz-SEP was examined. The results were largely replicated in the CCA-SEP: For the N80 component, ANOVA revealed a significant main effect of postural task (F_3,64_ = 24.688, p < 0.001, partial η^2^ = 0.536), with significant differences in the post hoc comparisons. For the P40 component, ANOVA also revealed a significant main effect of postural task (F_3,64_ = 9.802, p < 0.001, partial η^2^ = 0.314) and the same results of post hoc comparisons, except for the comparison between the QS and the CO.

### Comparison of PSD in prestimulus waveform: Cz vs. CCA

The main effect of Type was significant (α; F_1,112_ = 37.273, p<0.001; β; F_1,112_ = 14.238, p<0.001), indicating greater relative oscillatory power at Cz than at CCA. The distinct spectral characteristics provided an additional rationale for quantifying prestimulus amplitude from electrode-level EEG.

**Supplementary Figure 2.**
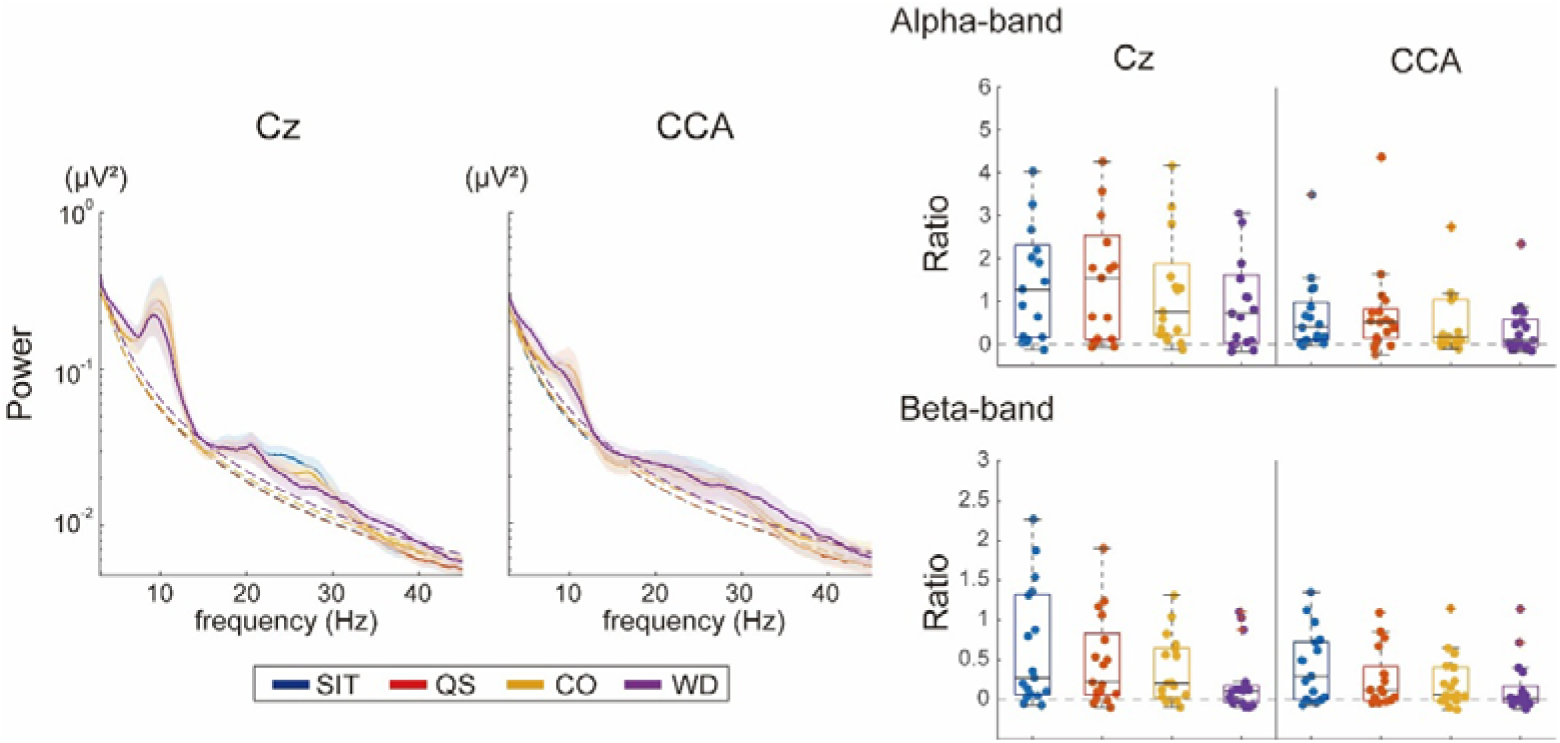
Prestimulus power spectral density and aperiodic-adjusted relative power. Prestimulus oscillatory amplitude was calculated in electrode space rather than the CCA waveform. To compare the oscillatory components in the α- and β-bands between the representative electrode, Cz, and CCA, power spectral density was computed using the FOOOF toolbox (Donoghue et al., 2020). LME was used to examine the effects of Type (Cz vs. CCA), Condition, and their interaction on relative oscillatory power (Ratio): (PSD original – PSD aperiodic) / PSD aperiodic.

Donoghue T, Haller M, Peterson EJ, Varma P, Sebastian P, Gao R, Noto T, Lara AH, Wallis JD, Knight RT, Shestyuk A & Voytek B. (2020). Parameterizing neural power spectra into periodic and aperiodic components. *Nat Neurosci* **23,** 1655-1665.

### Influence of different electrical stimulation on SEP: pulse-train stimulation vs. single-pulse stimulation

**Supplementary Figure 3.**
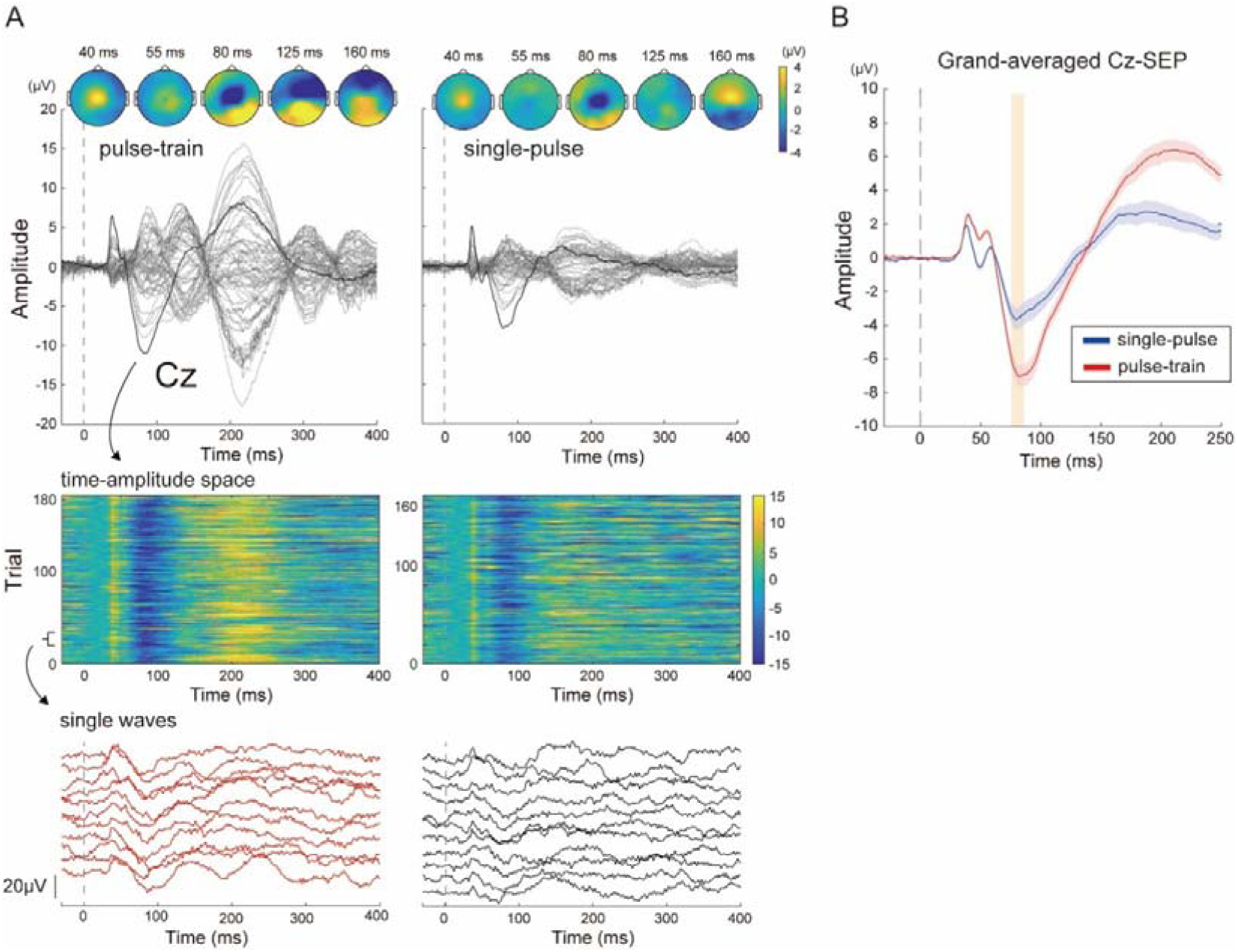
Similarity between SEP waveforms evoked by pulse-train stimulation and single-pulse stimulation. Compared with conventional single-pulse stimulation, we examined the characteristics of SEPs evoked by the pulse-train stimulation. In thirteen participants (27.3±2.1 years old, 168.3±1.9 cm; one participant was finally excluded from the analysis because of an error in the stimulus intensity setting), 200 additional SEPs were recorded using single-pulse stimulation (pulse duration: 1.0 ms) in the SIT condition. The intensity of single-pulse stimulation was adjusted so that its intensity relative to the individual stimulation threshold was equivalent to that used for pulse-train stimulation. For the recorded SEPs, we quantified the similarity between averaged waveforms evoked by single-pulse and pulse-train stimulations within each participant. We also quantified the similarity of each single-trial waveform to the corresponding averaged waveform that was calculated using a leave-one-out method for each stimulation procedure. Pearson’s correlation coefficients (r) were calculated over the 30-120 ms time window, and the r values were Fisher z-transformed in both analyses. The average SEP waveforms evoked by the two stimulation procedures showed high similarity (mean: r = 0.895, 95%CI: 0.812-0.943). For single-trial waveform similarity, the Fisher z-transformed correlation coefficients were averaged across trials within each participant and simulation procedure. A paired t-test revealed significantly higher similarity for pulse-train stimulation than for single-pulse stimulation [t(11) = 4.50, p<0.001, dz = 1.30, mean r = 0.774 (95%CI: 0.633∼0.866) vs. mean r = 0.561 (95%CI: 0.440∼0.662)]. Thus, pulse-train stimulation preserves the characteristics of the SEP waveform while improving the detectability of the response at the single-trial level. A. SEP waveforms from a representative single participant for pulse-train stimulation (left) and single-pulse stimulation (right). *Upper:* Average waveform across 64 EEG electrodes and corresponding scalp topographies. *Middle:* trial-to-trial EEG amplitudes over time. *Lower:* Representative single-trial waveforms. B. Grand-averaged waveforms of Cz-SEP across the twelve participants for pulse-train and single-pulse stimulation.

### Sensitivity analyses of the effects of prestimulus β-amplitude and COPv

To further assess the robustness of the association between prestimulus β-amplitude and CCA-SEP amplitude after accounting for postural sway, fixed-effect inference was evaluated using small-sample-corrected cluster-robust standard errors, with participant as the clustering variable. In addition, the model including COPv was compared with the corresponding model without COPv using a likelihood-ratio test.

The cluster-robust sensitivity analysis showed that the effect of prestimulus β-amplitude remained significant (b = 0.050, t_15.76_ = 3.984, p = 0.001), whereas the effect of COPv did not reach statistical significance (b = −0.032, t_14.64_ = −1.996, p = 0.064). Model comparison further showed that including COPv did not significantly improve model fit relative to the model without COPv (LRT: χ²(4) = 6.001, p = 0.199). Thus, both the cluster-robust sensitivity analysis and model comparison supported the robustness of the β-amplitude effect, whereas the independent contribution of COPv was less robust.

